# Single Cell Cytometry of Mouse Brain Tissue

**DOI:** 10.64898/2026.09.22.753505

**Authors:** Caroline E. Roe, Chiara Miller, Nidia Calero, Jonathan M. Irish, Andra L. Dingman

## Abstract

Suspension mass cytometry can measure 50 or more markers per cell and is not impacted by cellular autofluorescence, making it an appealing platform for comprehensive phenotyping of single cells in tissue with high innate fluorescence, such as brain. However, this organ is challenging to reduce to a single cell suspension: it is a deeply interconnected tissue rich in extracellular material and contains a wide variety of cell types including neurons, their supporting glia, and hematopoietic cells from peripheral circulation, any or all of which may have competing requirements for survival through the dissociation process.

This study compares approaches for producing single-cell preparations from white and gray matter–enriched regions of adult mouse brain, testing simple mechanical dissociation, Accutase, and three gentle enzymes plus DNase: papain, collagenase II, and collagenase plus dispase, all with or without Percoll gradient centrifugation to separate intact cells from debris. Live cells were counted by hemocytometer and Trypan blue exclusion to assess yield and compare across enzyme conditions. Mass cytometry was used to assess cellular composition after representative dissociation, debris removal, and viable cryopreservation. Key proteins measured included βIII-tubulin for neurons, glial fibrillary acidic protein (GFAP) for astrocytes, oligodendrocyte transcription factor (OLIG2) for oligodendrocytes, CD31 for endothelial cells, CD24 for ependymal cells, purinergic receptor P2Y12 (P2RY12) for microglia, and CD45 for leukocytes.

Mechanical disaggregation paired with enzymatic dissociation by papain, Collagenase/Disapse, or Collagenase II plus DNase produced approximately 400,000 live cells per 100 grams of tissue. Cell yield was not significantly different between conditions and yields were similar between white and grey matter. Addition of Percoll gradient centrifugation substantially reduced the number of viable cells recovered (average decrease of 88%) and resulted in viable cell numbers that were insufficient for robust analysis of less common cell subsets. In addition to neurons, mass cytometry identified six major populations of non-neuronal cells: oligodendrocytes, astrocytes, endothelial cells, ependymal cells, microglia, and leukocytes. The relative abundance of these six populations was generally unchanged by cryopreservation in both grey matter (*r* = 0.95), and white matter (*r* = 0.90), though loss of oligodendrocytes did occur in both tissues. Proliferation, as measured by Ki67 positivity, was non-existent; consistent with this, levels of phosphorylated STAT1 (p-STAT1) and p-STAT3 proteins were low in all cell populations and unchanged by cryopreservation. A modest increase in p-S6(S235/236) after cryopreservation was consistently observed only in microglia.

Together, these results establish standardized procedures for generating viable single cell suspensions for cytometry that preserve the cellular diversity of the adult murine brain with minimal perturbation of intracellular signaling cascades.

## Introduction

When preparing single cell suspensions from tissue for high parameter analysis, a primary goal is to maximize viable cell yield while maintaining the cellular diversity of the source material^1^. This is especially challenging in the mammalian brain, a matrixed organ composed of neurons sheathed in lipid-rich myelin as well as support cells known as glia, blood vessels, and immune cells of hematopoietic origin^2^. When disaggregated, this tissue forms a suspension of cells of varying size and morphology along with cellular and extracellular debris^3,4^. For such mixtures, most automated platforms tend to overcount cells, making manual counting by hemocytometer the preferred, if time-consuming, method to count live cells^5^. Additionally, neural tissues tend to have high but variable autofluorescence which challenges assays reliant on light for their measurements even with computational correction^6,7^. Fortunately, mass cytometry enables high-dimensional, per cell measurements without the need for fluorescent tags or light of any kind^8^. Mass cytometry has been widely used for deep phenotyping of a wide range of mammalian tissues^9^ including but not limited to human tonsil^10^, bone marrow^11^, ovary^12^, brain^13^, skin^14^, and mouse bone marrow^15^, spleen, gut^,16^ *s*keletal muscle^17^, kidney, heart, liver^18^, and brain^19^.

While numerous protocols for dissociation of mouse brain tissue are available, many focus on the survival of only one cell type of interest and are less concerned with absolute cell yield, overall viability, or the retention of all cell types in the tissue^20,21^. These protocols typically use combined mechanical and enzymatic dissociation^22^ and often use papain^23,24^, collagenase alone^25^, or collagenase plus dispase^26^. Propriety dissociation kits and systems such as GentleMACs are also available ^27,28^. To reduce the quantity of confounding debris in the sample, many protocols suggest a debris removal step, such as Percoll gradient centrifugation^29,30^. However, depending on the type of gradient and density of Percoll solution or solutions used, this step may intentionally or inadvertently enrich the sample one cell type^31,32^. This may be why reviews of protocol efficacy for specific cell types exist, but the field lacks a systematic comparison of common conditions utilizing widely accessible reagents and a global assessment of cell types yielded from dissociation^33,34^.

In this study, mechanical and enzymatic dissociation protocols were systematically tested on both white and grey brain matter from adult mice to identify an efficient, reliable method for dissociation of the entire mouse brain. Brain cells are differently sensitive to the conditions of dissociation^35^ so several conditions were compared for viable cells yield. Gentle enzyme conditions papain and collagenase plus dispase were identified in the literature and collagenase II was chosen based on prior work in human tissue, which also informed inclusion of DNase I in these three conditions^36^. Accutase was included as a stronger alternative to the above conditions. This enzyme mixture is commonly used to passage similar cells in culture as well as digest heavily myelinated tissue such as spinal cord^37^ or rat striatum^38^. Mechanical-only dissociation was included as reference points for cell yield without enzymes. Commercially available kits were avoided, in the interest of developing a transparent, economical protocol accessible to most researchers with access to fresh mouse brain tissue.

Due to tissue volume and cell number constraints, collagenase plus dispase was selected as the sole condition for tissue deep phenotyping by mass cytometry. This condition was selected because it performed as well as any other gentle enzyme in the prior comparison, is common in the literature^39^ and likely compatible with surface antibody staining^40^. Mass cytometry assessed the phenotype and abundance of major brain cell types, including neurons, astrocytes, oligodendrocytes, ependymal cells, and microglia, as well as hematopoietic and endothelial cells^41^. Abundances were calculated in freshly dissociated cells, the same cells after Percoll gradient centrifugation or after viable cryopreservation and thaw to determine the impact, if any, of these additional protocol steps to cell yield and quality.

## Results

### Gentle enzyme conditions tested have similar cell yields

Three gentle enzymatic dissociation conditions with DNase (papain, collagenase II, and collagenase plus dispase) were tested alongside mechanical dissociation alone and Accutase digestion to identify optimal conditions for viable cell yield from mouse brain white and grey matter as shown in **Figure 1**. Cell yield per gram was consistent with yield from human gliomas in published protocols^42^ and was approximately one third the literature estimated number of cells per mm^3^ based on imaging studies^43^. There was no statistically significant difference in viable cell yield between gentle dissociation conditions without Percoll gradient centrifugation in either white or grey matter, and these three conditions were not significantly different from mechanical only dissociation (all *P* > 0.05). Accutase treatment was superior to all three gentle enzyme conditions in yielding live cells from white matter only (*P* = 0.02).

**Figure 1.**
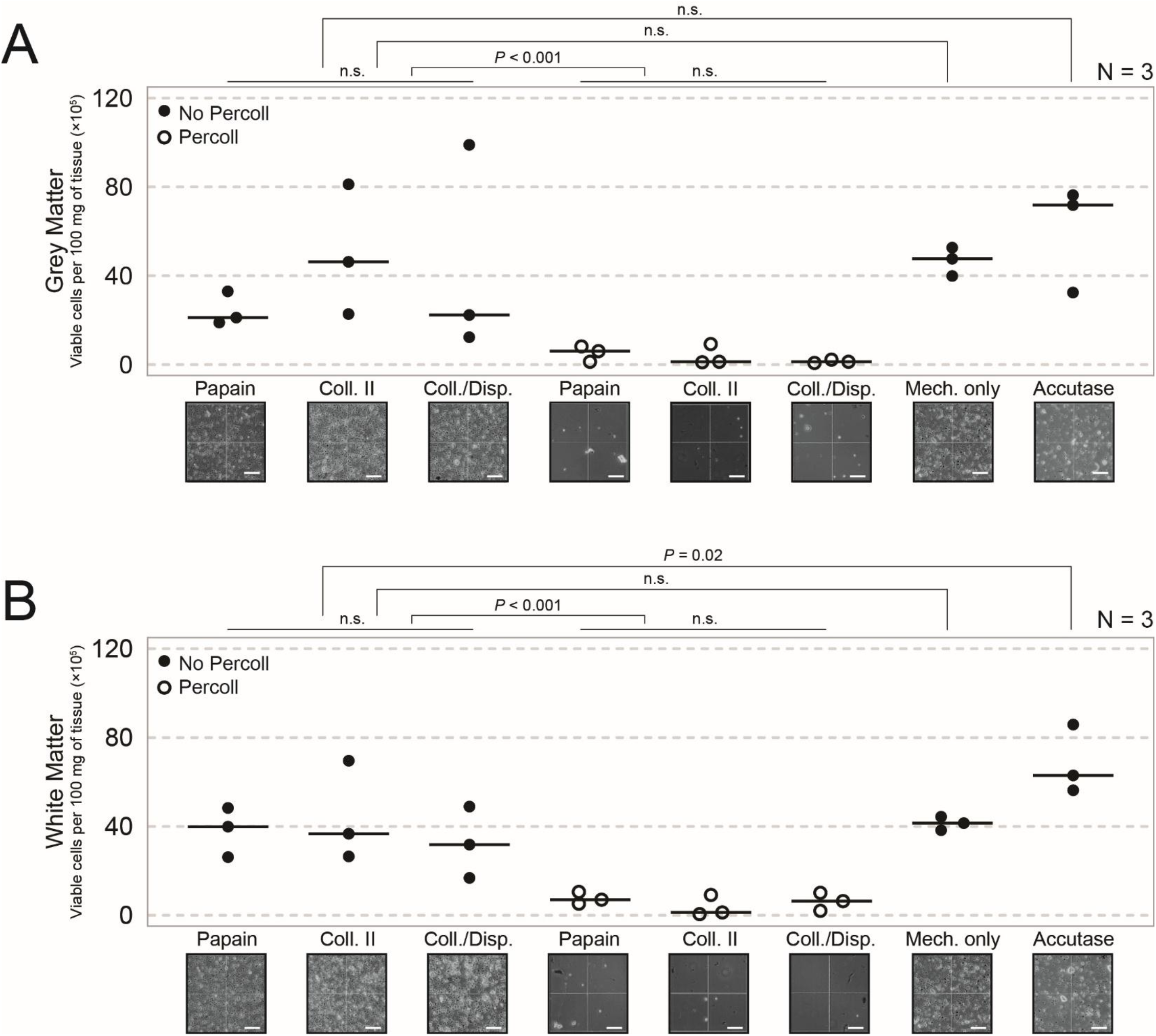
Enzymatic digestion without Percoll gradient centrifugation yields live cells from mouse brain. Graphs show yield across tissue dissociation conditions as hundreds of thousands of viable cells per 100 grams, in both mouse gray matter (A) and white matter (B). Each dot indicates a technical replicate (N = 3), and thick horizontal line indicates mean yield per condition. Conditions left to right: Mech. Only: mechanical only dissociation without enzymes, Accutase: 10’ Accutase digestion without other enzymes, then three gentle enzyme conditions with DNase: Papain: 30 minute papain digestion, Coll. II: 30 minute collagenase II digestion, Coll./Disp.: 30 minute collagenase plus dispase digestion. Yield for gentle enzyme conditions was calculated before (black dots) and after (white dots) Percoll gradient centrifugation to remove debris. Representative Trypan blue stained images are under each condition. Scale bars = 100µm. Statistically significant comparisons are denoted by p-values, p-values > 0.05 are indicated by n.s.

### Percoll gradient centrifugation reduces yield and skews sample composition

A 22% Percoll gradient was used to remove myelin and cell debris from dissociated tissue suspensions. A large quantity of material was removed greatly facilitating cell counting, as shown in **Supplemental Figure 1**, but viable cell yields after Percoll gradient centrifugation were significantly diminished for both white and grey matter (*P* < 0.001), with an average yield of 13.34% across three technical replicates, three enzyme conditions, and two tissue types (range = 1.79%-28.72%). This cell loss was compounded when a Percoll step was included prior to staining and collection for mass cytometry, with 0.001% as many intact cells collected at the cytometer compared to those samples prepared without a Percoll step (mean_Fresh_ = 516,451 cells per sample, mean_Percoll_ = 552 cells per sample). Further, the composition of samples post-Percoll only partly resembled the same sample without Percoll (*R*_*grey*_ = -0.45, *R*_*white*_ *=* -0.78), with a consistent loss of oligodendrocytes and ependymal cells across tissue types and a commiserate enrichment in leukocytes, microglia, and endothelial cells, reported in **Table 1**. Overall reduction in non-neuronal cells precluded deeper analysis of individual cell subsets from samples, though absolute cell counts for these populations are reported in **Supplemental Table 1**.

**Table 1.** Event counts and frequency of terminal populations in freshly prepared white and grey matter and the same tissue after Percoll gradient centrifugation. Percent abundance of each population was calculated as a percentage of all non-neuronal cells assigned to any listed population (Event count: Total). Correlation between population abundances before and after Percoll gradient centrifugation is reported with Pearson’s correlation coefficient *r* and associated *p*-value.

|  | Subsets | Event count |  | Percent |  |  | *r* |
| --- | --- | --- | --- | --- | --- | --- | --- |
|  |  | Fresh | Percoll | Fresh | Percoll | Delta |  |
| Grey matter | Oligodendrocytes | 542 | 0 | 15.4 | 0 | -15.4 | -0.45<br>*P* = 0.38 |
|  | Astrocytes | 1268 | 3 | 35.9 | 12.5 | -23.4 |  |
|  | Microglia | 130 | 3 | 3.7 | 12.5 | +8.8 |  |
|  | Leukocytes | 136 | 14 | 3.9 | 58.3 | +54.4 |  |
|  | Ependymal cells | 1272 | 1 | 36 | 4.2 | -31.8 |  |
|  | Endothelial cells | 181 | 3 | 5.1 | 12.5 | +7.4 |  |
|  | **Total** | **3529** | **24** |  |  |  |  |
| White matter | Oligodendrocytes | 1626 | 2 | 27.4 | 6.1 | -21.3 | -0.78<br>*P* = 0.678 |
|  | Astrocytes | 698 | 8 | 11.8 | 24.2 | +12.4 |  |
|  | Microglia | 213 | 5 | 3.6 | 15.2 | +11.6 |  |
|  | Leukocytes | 583 | 10 | 9.8 | 30.3 | +20.5 |  |
|  | Ependymal cells | 2528 | 1 | 42.6 | 3 | -39.6 |  |
|  | Endothelial cells | 287 | 7 | 4.8 | 21.2 | +16.4 |  |
|  | **Total** | **5935** | **33** |  |  |  |  |

### Dissociation of mouse brain by collagenase II plus dispase without Percoll yields glia

Expert gating, as depicted in **Figure 2A**, identified seven major cell subsets expected in this tissue^44^. In the order they were identified, these subsets were: β-III tubulin+ neurons^45^, OLIG2+ oligodendrocytes^46^, GFAP+ astrocytes^47^, P2RY12+ brain resident microglia^48^, CD45hi blood-derived leukocytes^49^, CD31+ endothelial cells^50^, and CD24+ ependymal cells^51^. These seven populations were present in all tissues and conditions analyzed. Across all samples, β-III tubulin+ neurons were the most abundant cell type, consistent with imaging assessments and other single-cell studies of adult mouse brain^52,53^. Further gating for immune cell subsets in these conditions identified three populations of hematopoietic cells in all samples: NK1.1+ natural killer cells^54^, CD11b-blood-derived monocytes, and CD11b+ blood-derived macrophages^55^. In the context of this manuscript, we will use the term blood-derived macrophage to refer to cells of the myeloid lineage originating in the bone marrow from the adult hematopoietic stem cell and common myeloid progenitor. We would conceptually distinguish these HSC-derived blood macrophages from tissue resident macrophages like brain microglia that are thought to originate from embryonic yolk sac cells. CD3e+ T cells^56^ and B220+ B cells^57^ were also identified with this panel but were not detected in every sample. Further characterization of glia in **Figure 2B** demonstrated the purity of oligodendrocyte and astrocyte populations. **Figure 2C** confirmed co-expression of Ly6-C with CD31 in endothelial cells, and the absence of either of these targets on putative ependymal cells^50^. Deep phenotyping of microglia, blood-derived macrophages, and monocytes in **Figure 2D** confirmed low expression of CD45 on microglia, absence of CD11b on monocytes, and CX3CR1 expression by all three subsets^58^. CD44 was detected on monocytes, but not macrophage subsets^59^. LyC was absent from monocytes in this dataset, suggesting a non-classical, patrolling phenotype^60^.

**Figure 2.**
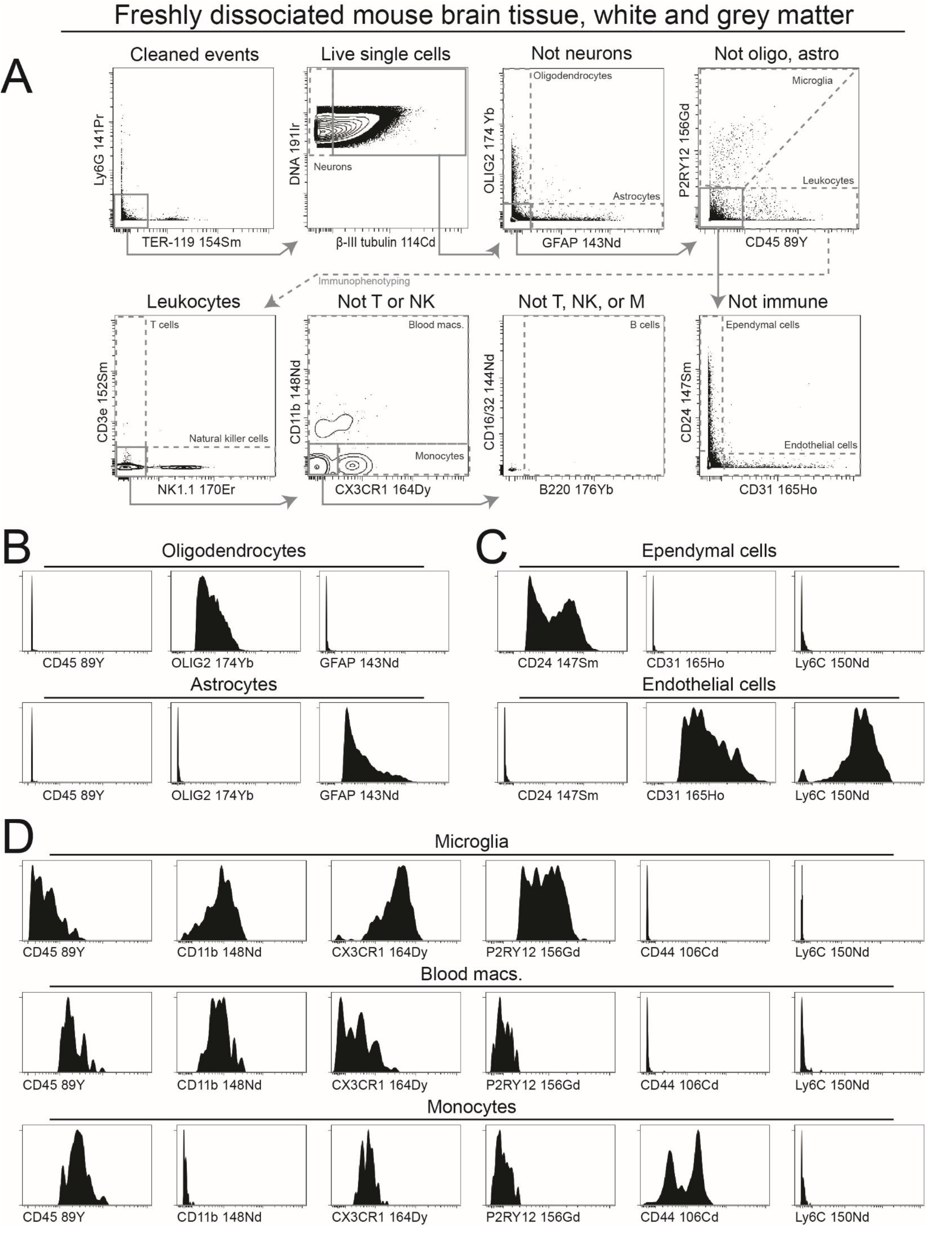
Mouse brain antibody panel identifies seven major cell subsets and five leukocyte subsets. Biaxial plots show gating for established cell types in mouse brain tissue prepared using collagenase/dispase dissociation protocol. (A) From live intact single cells, TER-119+ red blood cell and Ly6G+ granulocyte events are excluded. This is followed by sequential gating of β-III tubulin+ neurons, OLIG2+ oligodendrocytes, GFAP+ astrocytes, CD31+ endothelial cells, P2RY12+ microglia, CD45hi leukocytes, CD24+ ependymal cells and CD31+ endothelial cells. Frequency of terminal populations (dashed line gates) was assessed under relevant protocol steps in Tables 2 and 3. Major populations of blood-derived CD45hi leukocytes are gated: CD3e+T cells, NK1.1+ natural killer cells, CD11b-monocytes and CD11b+ blood-derived macrophages, and B220+ B cells. (B) Comparison of oligodendrocyte and astrocyte phenotypes. (C) Comparison of ependymal cell and endothelial cell phenotypes. (D) Deep phenotyping of microglia, myeloid-derived macrophages, and monocytes.

### Mouse brain and immune cell subsets are stable through cryopreservation

High correlation between cell frequencies in the sample samples before and after thaw (*R*_*grey*_ = 0.95, *R*_*white*_ *=* 0.90) as reported in **Table 2** indicated that cryopreservation had no significant impact on sample composition. Despite concerns about changes in marker expression due to stress from freeze/thaw^61^, phenotypic analysis of these subsets detected no major changes in any identity measurements for individual subsets, either major or immune, after cryopreservation, as depicted in **Figure 3**. No Ki67+ proliferating cells were observed in any sample from this study, shown in **Figure 4A**. Levels of p-STAT1 and p-STAT3 were generally low and unchanged by cryopreservation, in **Figure 4B**. p-S6 (S235/236), was slightly elevated only in microglia after cryopreservation and thaw, shown in **Figure 4B** and **Figure 4C**, arcsinhFC_white_ = 0.16, arcsinhFC_grey_ = 0.06.

**Table 2.** Event counts and frequency of terminal populations in freshly prepared white and grey matter and the same tissue after cryopreservation and thaw. Percent abundance of each population was calculated as a percentage of all non-neuronal cells assigned to any listed population (Event count: Total). Correlation between population abundances before and after cryopreservation is reported with Pearson’s correlation coefficient *r* and associated *p*-value.

|  | Subsets | Event count |  | Percent |  |  | *r* |
| --- | --- | --- | --- | --- | --- | --- | --- |
|  |  | Fresh | Cryo | Fresh | Cryo | Delta |  |
| Grey matter | Oligodendrocytes | 542 | 99 | 15.4 | 4.2 | -11.2 | 0.95<br>*P* < 0.05 |
|  | Astrocytes | 1268 | 956 | 35.9 | 41 | +5.1 |  |
|  | Microglia | 130 | 113 | 3.7 | 4.8 | +1.1 |  |
|  | Leukocytes | 136 | 69 | 3.9 | 3 | -0.9 |  |
|  | Ependymal cells | 1272 | 881 | 36 | 37.8 | +1.8 |  |
|  | Endothelial cells | 181 | 215 | 5.1 | 9.2 | +4.1 |  |
|  | **Total** | **3529** | **2333** |  |  |  |  |
| White matter | Oligodendrocytes | 1626 | 363 | 27.4 | 15.8 | -11.6 | 0.90<br>*P* < 0.05 |
|  | Astrocytes | 698 | 181 | 11.8 | 7.9 | -3.9 |  |
|  | Microglia | 213 | 89 | 3.6 | 3.9 | +0.3 |  |
|  | Leukocytes | 583 | 85 | 9.8 | 3.7 | -6.1 |  |
|  | Ependymal cells | 2528 | 1341 | 42.6 | 58.5 | +15.9 |  |
|  | Endothelial cells | 287 | 232 | 4.8 | 10.1 | +5.3 |  |
|  | **Total** | **5935** | **2291** |  |  |  |  |

**Table 3.** Mass cytometry mouse brain phenotyping panel.

| Antigen | Tag | Clone | Vendor | Catalog # | RRID | Staining |
| --- | --- | --- | --- | --- | --- | --- |
| CD45 | 89Y | 30-F11 | Standard Biotoools | 3089005B | AB_2651152 | Live |
| CD44 | 106Cd | IM7 | Standard Biotoools | 92J005106 | AB_3678360 | Live |
| beta-III-tubulin | 114Cd | TUJ1 | Biolegend | 801201 | AB_2313773 | Post-perm |
| Ly-6G | 141Pr | 1A8 | Standard Biotoools | 3141008B | AB_2814678 | Live |
| CD11c | 142Nd | N418 | Standard Biotoools | 3142003B | AB_2814737 | Live |
| GFAP | 143Nd | GA5 | Standard Biotoools | 3143022B | AB_2938640 | Post-perm |
| CD16/32 | 144Nd | 93 | Standard Biotoools | 3144009B | AB_2814674 | Live |
| CD4 | 145Nd | RM4-5 | Standard Biotoools | 3145002B | AB_2687832 | Live |
| CD24 | 147Sm | MI/69 | eBiosciences | 50-124-83 | AB_467170 | Live |
| CD11b | 148Nd | MI/70 | Standard Biotoools | 3148003B | AB_2814738 | Live |
| Ly-6C | 150Nd | HK1.4 | Standard Biotoools | 3150010B | AB_2895118 | Live |
| CD3e | 152Sm | 145-2C11 | Standard Biotoools | 3152004B | AB_2687836 | Live |
| p-STAT1 (Y701) | 153Eu | 58D6 | Standard Biotoools | 3153003A | AB_2811248 | Post-perm |
| TER-119 | 154Sm | TER-119 | Standard Biotoools | 3154005B | AB_3677870 | Live |
| P2RY12 | 156Gd | S16007D | Biolegend | 848002 | AB_2650634 | Live |
| p-STAT3 (Y705) | 158Tb | 4-p-STAT3 | Standard Biotoools | 3158005A | AB_2811100 | Post-perm |
| TCR $\gamma/\delta$ | 159Tb | GL3 | Standard Biotoools | 3159012B | AB_2922919 | Live |
| Ki67 | 162Dy | B56 | Standard Biotoools | 3162012B | AB_2888928 | Post-perm |
| CX3CR1 | 164Dy | SA011F11 | Standard Biotoools | 3164023B | AB_2832247 | Live |
| CD31 | 165Ho | 390 | Standard Biotoools | 3165013B | AB_2801434 | Live |
| CD8a | 168Er | 53-6.7 | Standard Biotoools | 3168003B | AB_2811241 | Live |
| NK1.1 | 170Er | PK136 | Standard Biotoools | 3170002B | AB_2885023 | Live |
| CD86 | 172Yb | GL1 | Standard Biotoools | 3172016B | AB_2922923 | Live |
| OLIG2 | 174Yb | 211F1.1 | Sigma-Aldrich | MABN50 | AB_10807410 | Post-perm |
| p-S6 (S235/236) | 175Lu | N7-548 | Standard Biotoools | 3175009A | AB_2661838 | Post-perm |
| B220 | 176Yb | RA3-6B2 | Standard Biotoools | 3176002B | AB_2895123 | Live |

**Figure 3.**
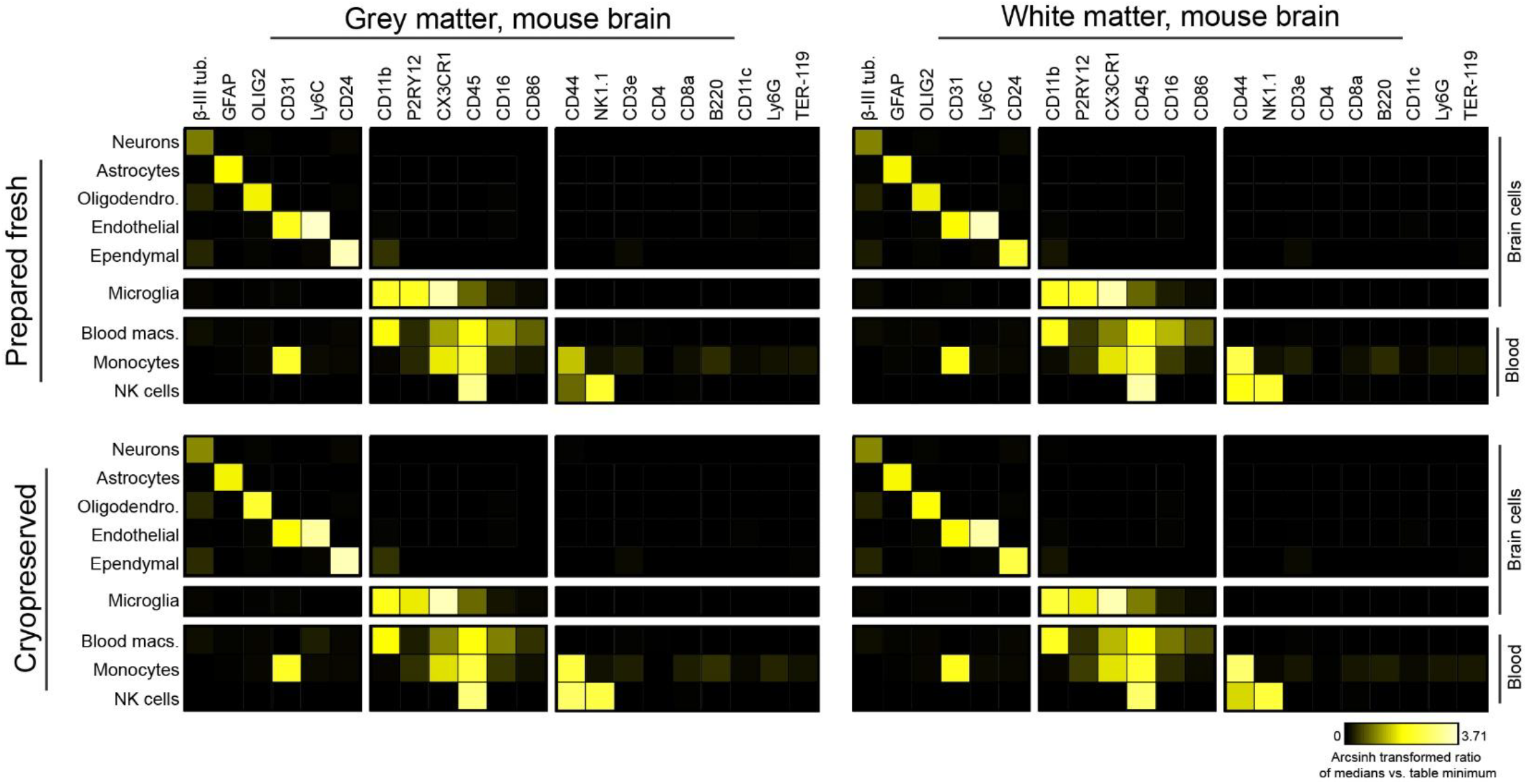
Phenotype of major subsets is stable through cryopreservation. Heatmaps show arcsinh-transformed relative median fold change of mass intensity versus table minimum (0) for phenotypic markers across identified cell subsets. Heatmap blocks are divided by cells of neural origin (Brain cells) and adult hematopoietic origin (Blood) with microglia separated to indicate their unique function as a brain-derived immune cell. Columns of markers are likewise divided into three categories (L to R) general neural markers, macrophage markers, and all other immune targets.

**Figure 4.**
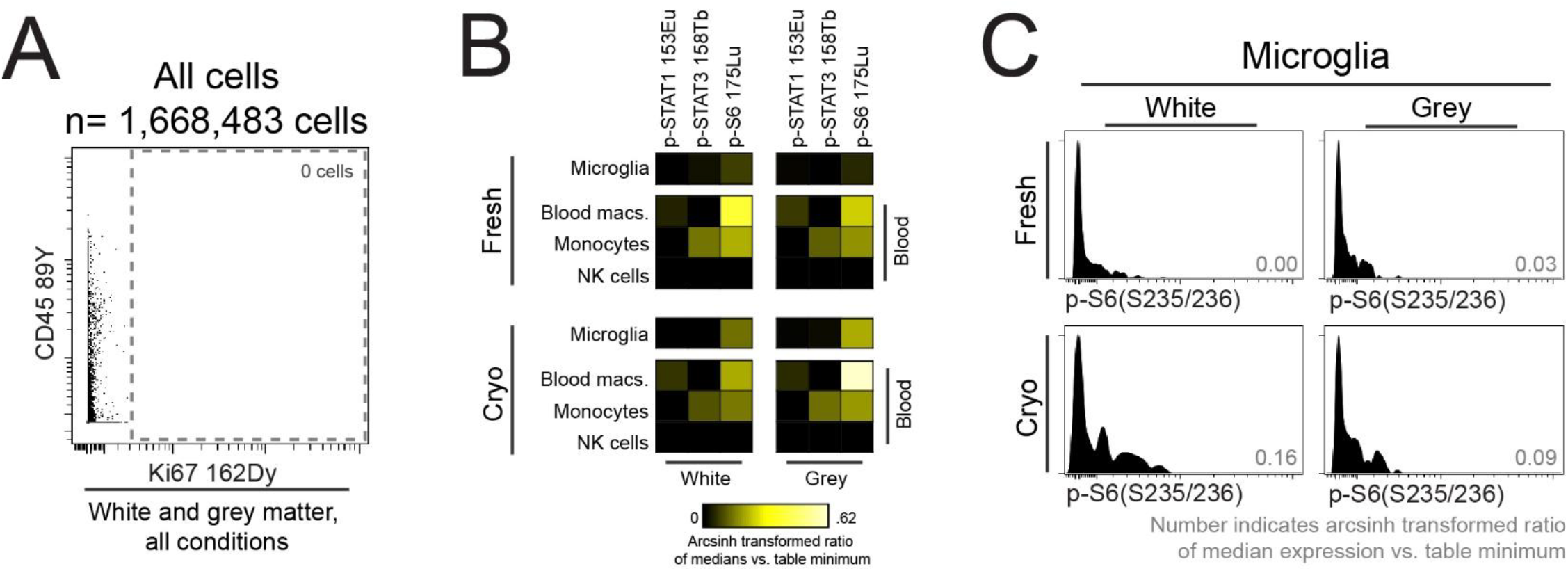
No proliferation, but some signaling is observed in dissociated mouse brain cells. In part A, no Ki67+ non-proliferating cells are present in any preparation condition or tissue. Plot is of all pooled live intact cells from all conditions (fresh, post-Percoll, and cryopreserved) and all tissue (white and gray matter). In part B, some levels of basal p-STAT1, p-STAT3, and p-S6 (S235/236) is present in microglia and myeloid lineage cell subsets. The only signaling pattern changed by cryopreservation is shown in C, a slight increase in p-S6 (S235/236) in microglia from both tissues.

## Discussion

Gentle enzymatic dissociation with DNase and collagenase plus dispase yielded on average 385,000 live cells per 100 milligrams of mouse brain tissue and provided cells of sufficient quality for mass cytometry phenotyping of all cell types expected in the tissue. Papain, collagenase, and collagenase/dispase are comparable to each other as well as to harsher digestion with Accutase and enzyme-free mechanical dissociation. As enzymatic digestion can cleave extracellular epitopes it is suggested to use the gentlest, most specific, conditions for flow cytometry assays to maximize the likelihood of preserving surface epitopes of interest and ensure survival of cells through the dissociation process^62^. Accutase can kill astrocytes in neural cell suspensions^63^. Papain is a broad cysteine protease that has long been known to cleave a variety of extracellular targets in a manner that may inhibit detection of these targets^64^. In contrast, collagenase plus dispase targets specifically collagen and fibronectin^65^, likely sparing surface epitopes. While staining of key markers in this study looked as expected after collagenase plus dispase exposure, rigorous testing of the impact of enzyme selection on surface antibody staining is required to identify the optimal enzymes for dissociation of this tissue.

Percoll gradient centrifugation, used for removal of myelin and cellular debris, reduced cell yield from dissociation by almost 90%. This reduction in yield, coupled with the small tissue volumes available from individual mouse brains, makes downstream assays requiring millions of cells, such mass cytometry, extremely challenging^66^. Multiple samples can be pooled, but mouse genotype, brain region of interest, time, and cost may make this impractical. For downstream assays which are sensitive to on-cellular material, such as single-cell RNA sequencing, this reduction in total cell number may be an acceptable tradeoff^67^. If debris is not removed physically or by a threshold set at the instrument, cytometry technologies that do not detect unmarked non-cellular material, such as mass cytometry, may perform better than traditional or spectral flow cytometry^68^. Detection methodology can minimize the impact of debris; for example, this study used iridium-labelled DNA intercalator rather than cisplatin for cell detection to avoid inadvertent labelling of protein aggregates or other debris in tissue suspensions^69^.

Encouragingly, viable cryopreservation appeared to have minimal effect on the abundance and phenotype of major cell types. The abundance of major populations, as well as immune subsets, are highly correlated between fresh and cryopreserved samples and align with cell types identified in other studies and by other single-cell technologies. No population analyzed here was completely lost due to cryopreservation. Proliferation, as measured by Ki67+ cells, was non-existent in all samples in this study. While proliferation is thought to be extremely rare in the uninjured, healthy adult brain, some studies have provided evidence for it^70^. It is possible that a single proliferation maker is insufficient to detect all proliferating cells, and other measurements of cell cycle progression, such as proligera proliferating cell nuclear antigen (PCNA) or 5-Iodo-2′-Deoxyuridine (IdU), could be valuable, if accurate assessment of proliferation is essentia^l71,72^. Baseline cell signaling, measured at three selected phospho-proteins, was comparable before and after cryopreservation except for a modest increase in p-S6-S235/236 specifically in microglia from white or grey-matter. This finding suggests that microglia may be activated in part by some aspect of the freeze/thaw process that does not affect any of the other cell types. Longer rest periods after thaw could allow activated cells to return to baseline. For functional studies of mouse brain cells, cryopreservation should be evaluated on a readout-by-readout and cell-by-cell basis to document the impact, if any, on the dynamic measurement of interest.

Overall, this work evaluates key preparation steps of suspended mouse brain tissue for high dimensional antibody-based protein measurements by suspension flow cytometry. These data provide important options for investigators based on experimental goals. Enzymatic digestion yielded high viable cell numbers that recapitulated cell types *in vivo*. This approach allows for simultaneous evaluation of protein expression and intracellular signaling among many CNS cell types. Debris removal has long been critical to creating single cell suspensions from mouse brain tissue for fluorescent cytometry and RNA sequencing, but here resulted in significant loss of viable cells. Our data also showed that cell types were differentially preserved with density gradient debris removal. A distinct advantage of mass cytometry over other types of flow cytometry is that the contribution to event counts can be minimized. Importantly, these data suggest that viably cryopreserved cells isolated with this protocol are generally comparable to their fresh counterparts and suitable for a variety of downstream assays that require viable cells. This is particularly important for longitudinal studies using samples collected over multiple time points, as it allows for sample storage and batch processing.

## Materials and Methods

### Mouse brain tissue collection

All animal procedures were carried out in accordance with institutional (Institutional Animal Care and Use Committee) and National Institute of Health guidelines. Mice were wild-type C57/BL6 (MGI:2159769) females aged 60-120 days. Animals were anesthetized with isoflurane, transcardially perfused with cold PBS and the brain grossly resected into myelin-rich white matter regions, namely cerebellar white matter and brain stem, and more cellular grey matter portions, in this case cortex and hippocampus. Tissue was weighed prior to being placed in room temperature PBS for transport to the laboratory, with each animal yielding 150-200 milligrams of each tissue type. Tissue processing began within 60 minutes of animal sacrifice.

### Dissociation and cryopreservation reagents

Reagents used, with manufacturer, vendor, and catalog numbers, are reported in **Supplementary Table 2**. Serum-free media appropriate for mouse neural cells was used, namely DMEM/F12 with GlutaMAX and B27 Supplement. DNase I was used at a final concentration of 0.25 mg/mL in papain, collagenase II, and collagenase/dispase conditions. Papain was used at ∼20U/mL. Collagenase II and collagenase/dispase were used at 1 mg/mL. Accutase was used at half the manufacturer’s recommendation for tissue culture, 1:1 with serum-free media defined above. Cells were cryopreserved, when applicable, in the above media with 12% DMSO by volume, stored overnight at -80°C in a controlled cooling container before transfer to vapor-phase liquid nitrogen for storage until thaw. Upon thaw, cryopreserved cells rested in media plus DNase 1 for 20 minutes at 37°C before proceeding to antibody staining. 22% Percoll was prepared by diluting Percoll PLUS to 22% v/v in HBSS.

### Mechanical and enzymatic dissociation

For all conditions, tissue was finely minced with disposable #11 scalpels as previously described^73^ until in <1mm pieces, taking no longer than 5 minutes. To test mechanical dissociation alone, trituration and straining as described below occurred immediately after this mincing. Accutase condition was incubated 10’ at 37°C prior to mechanical dissociation. Papain, collagenase II, and collagenase plus dispase enzymatic dissociations included the addition of DNase I. These incubations were performed in a 37°C hybridization oven at low rotation for 30 minutes before mechanical dissociation. For all conditions, tissue was mechanically dissociated by trituration with a P1000 pipette until a homogenous, milky suspension was achieved, no longer than 5 minutes. The suspension was strained through 70 μm and 40 μm cell strainers prior to counting. For Percoll gradient centrifugation, cell suspensions were overlaid on 22% Percoll and centrifuged at 600×g for 15 minutes with no brake. Myelin layer and Percoll were gently removed and cells washed with HBSS before further use or counting. For comparison of enzymatic digestion conditions shown in **Figure 1**, tissue from two animals was pooled for coarse mincing before being split equally across all tested conditions per tissue type. For cytometry deep phenotyping, tissue from three animals was pooled and digested together with collagenase plus dispase, then split for comparison of freshly dissociated tissue versus after Percoll gradient reported in **Table 1** or viably cryopreserved dissociated tissue without Percoll gradient centrifugation, reported in **Table 2** and depicted in **Figure 2**, and **Figure 3**.

### Quantification of cell yield

Cells obtained from different dissociation protocols were manually counted on a hemocytometer using Trypan Blue exclusion to identify intact single cells^74^. Cell counts were normalized to the initial tissue weight and reported as tens of thousands of live cells per 100 mg of tissue.

### Mass cytometry

Antibody panel, reported in **Table 3**, was designed to identify all major brain cell types in the adult mouse, as well as identify major immune populations. The panel also included three signaling readouts: STAT1 (Y701) for interferon response, STAT3 (Y705) to detect response to IL-6, and the serine 235/236 phosphorylation site of ribosomal protein S6 to detect translation and protein synthesis^75,76^. Proliferation was measured by Ki67 positivity^77^. Metal-tagged antibody cocktails were prepared for the entire study, aliquoted and frozen as published and thawed as needed^78^. Cells were stained live for cell surface markers and viability^79^, fixed, permeabilized, stained for intracellular targets and total DNA according to established mass cytometry protocols^80,81^. Cells were stained with metal-tagged antibodies diluted as reported in **Table 3**, in 50 μL cell staining media for 30 minutes at room temperature. Mass cytometry data was collected on a CyTOF XT mass cytometer (Standard BioTools) at the CU Anschutz Cancer Center Flow Cytometry Shared Resource. Data was collected simultaneously on freshly dissociated tissue, the same tissue after Percoll density centrifugation to remove debris, and the same tissue after viable cryopreservation and thaw.

### Mass cytometry data analysis

Data were normalized with Standard BioTools bead-based normalization^82^ software prior to further analysis using Cytobank (RRID:SCR_014043)^83^. Data were manually gated in Cytobank to identify intact single cells alive at the time of antibody staining before expert gating for cell subsets. Gating scheme for defined brain cell types and immune cell subsets is depicted in **Figure 2**.

### Statistical testing

Cell counts from expert gated data were exported from Cytobank for statistical analysis in GraphPad prism (RRID:SCR_002798). Cell yield between tissue types was assessed using an unpaired t-test. The three gentle enzyme dissociation conditions with or without Percoll gradient centrifugation were compared as groups using Kruskal-Wallis tests. The change in cell abundances after Percoll gradient centrifugation was assessed by a Mann Whitney test per tissue type. Mechanical dissociation alone or Accutase digestion were compared to gentle enzyme conditions without Percoll using unpaired t-tests.

## Data Availability

Annotated mass cytometry data analyzed in this manuscript are publicly available online at Zenodo: https://doi.org/10.5281/zenodo.22831948.

## Author Contributions

CER, JMI, and ALD designed and conceptualized the study. NC and ALD provided freshly resected tissue specimens. CER and CM carried out protocols on tissue and collected data. CER, NC, JMI, and ALD interpreted data. CER, JMI, and ALD wrote the manuscript. JMI provided financial support. All authors contributed to reviewing and editing the manuscript.

## Acknowledgements

We thank the CU Flow Cytometry Shared Resource at the Cancer Center and the Human Immune Monitoring Shared Resource for reagent assistance. Special thanks to Dr. Rebecca A. Ihrie for providing advice and expertise regarding neural cell suspensions from primary mouse brain.

