## Supplemental Figure 1 for "Single Cell Cytometry of Mouse Brain Tissue"

Single Cell Cytometry of Mouse Brain Tissue
Roe et al. Supplemental Figure 1

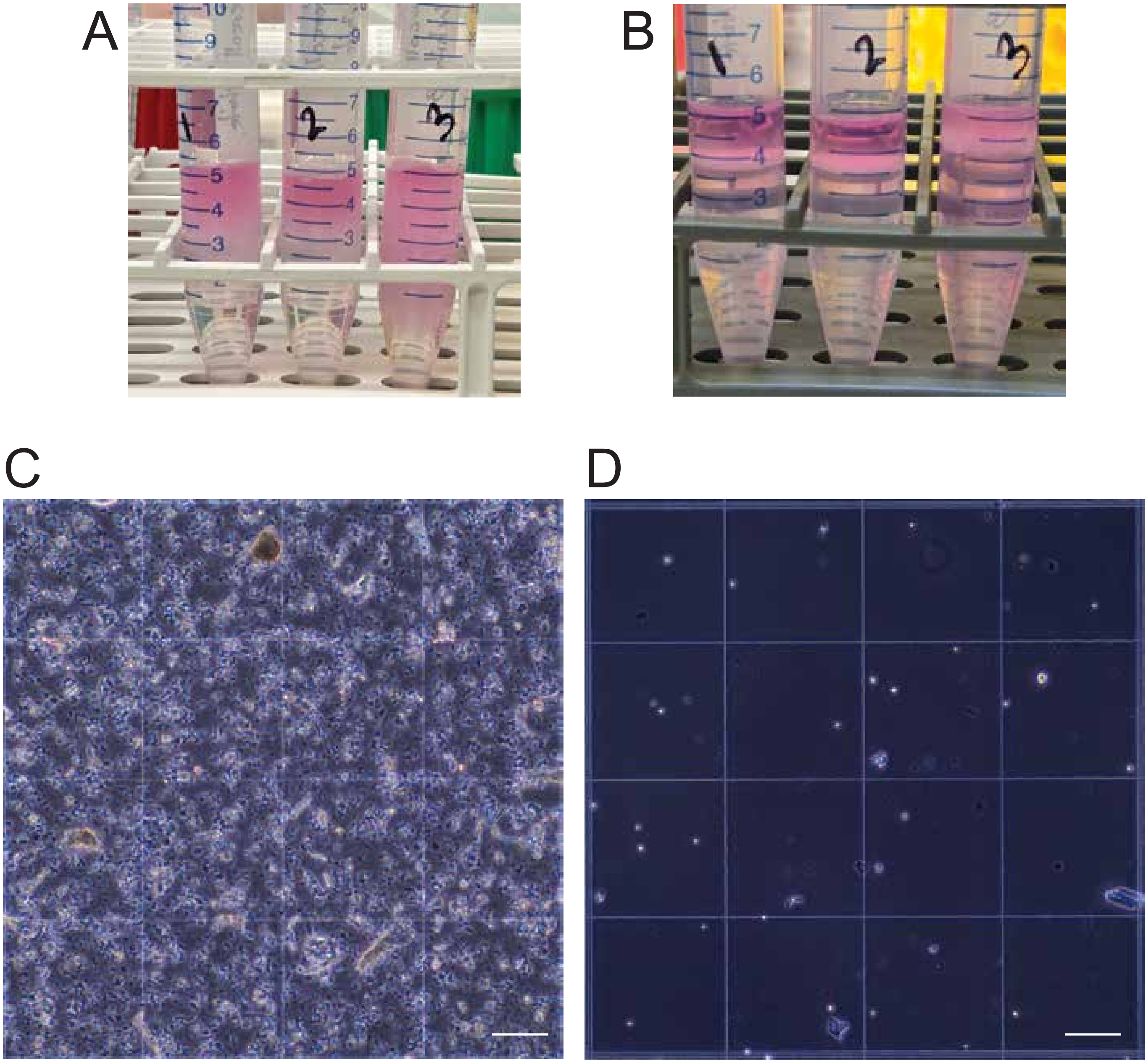


**Supplemental Figure 1 - Percoll gradient centrifugation removes most debris but also cells**
In A, cell suspension in pink media is layered on top of clear 22% Percoll.
After centrifugation,pink media is seperated from clear Percoll by a white layer of myelin and debris (B). Layers are removed and cell pellet at the bottom of the tube (not shown) is retrieved.
C and D are representative hemocytometer images of cells from A and B, respectively.
Before cleanup (C), abundant debris and cell clumps are present though individual live cells can be seen (n=73). After Percoll cleanup (D) retrieved cells are fewer (n=26) but suspension is almodevoid of debris and cell clumps, and thus easier to visualize.
